# Prototype matching: children’s preference for forming scientific concepts

**DOI:** 10.1101/2022.11.28.518150

**Authors:** Zhong Wang, Yi Zhang, Yi Jiang

## Abstract

Inspired by a sample lesson, this paper studies and discusses children’s preferences in learning scientific concepts. In a “Dissolution” lesson, one of the students took the demonstration experiment of “carmine dissolves in Water” demonstrated by the teacher as the prototype to judge whether a new phenomenon belongs to dissolution, instead of analyzing and judging the phenomenon by using the dissolution definition. Therefore, we propose a conjecture that “prototype matching” may be a more preferred way for children to learn concepts than thinking through inquiry experiment, analysis, deduction, etc. To this end, we conducted a targeted test on 160 fifth grade students (all of whom had learned this lesson) from a primary school in Beijing, and used goodness of fit test to statistically analyze the results. The results showed that: ➀ the Chi square of the general result is 73.865, P<0.001, indicating that children did have obvious prototype preference; ➁ We “tampered” some of the prototypes, that is, they looked like the prototypes that the teacher had told students, but they were actually wrong. However, the results showed that children still preferred these so-called “prototypes” (chi square is 21.823, P<0.001). Conclusion: ➀ Children have an obvious preference for “prototype matching” in scientific concept learning, which is not only obviously deviated from the current general understanding of science education that emphasizes inquiry construction, but also points out that there may be a priority relationship among various ways of concept organization (such as definition theory, prototype theory, schema theory, etc.). ➁ Children’s preference for prototypes seems to be unthinking, and they will not identify the authenticity of prototypes, which is particularly noteworthy in front-line teaching.

## 1. Introduction

This research originated from a science lesson of “dissolution”. In this lesson, the teacher asked the students to guess what factors might accelerate the dissolution. One of the girls replied that putting ice in hot water is accelerated dissolution. However, from the definition of dissolution, the melting of ice in hot water obviously does not belong to dissolution, because the premise of dissolution must be that two different substances are mixed together (Baike.baidu, “DISSOLUTION”). So why does this student think that ice melts in hot water is dissolved? To this end, we conducted an after-class interview with this student (see “6. Appendix” of this article for details). We noticed the last video. In this video, we did not directly ask her why the phenomenon of ice immersion in hot water is dissolution, but imagined a phenomenon: mixing blue and yellow pigments together to become green. Does this phenomenon belong to dissolution? Through this question, we try to understand how the student’s understand the concept of “dissolution”, and then understand why this student has the above point of view. However, it was unexpected that this student’s explanation of this hypothetical (she thought it was dissolution) was a little strange: instead of explaining and analyzing this phenomenon, she matched it with an experiment demonstrated by the teacher in that lesson, namely, the phenomenon that carmine dissolved in water. She thinks that this phenomenon is very similar to the demonstration experiment. Since the carmine is dissolution, naturally, this phenomenon (blue mix yellow become green) should also be dissolution.

Although this answer does not point to the original question, it leaves us a clue: it seems that we can see how this student understands the concept of “dissolution”. If we understand this problem, the problem of ice and hot water will be solved. From this question and answer, we can clearly feel that the student’s learning about the phenomenon of “dissolution” seems to follow “prototype matching”, that is, if the carmine experiment is known in advance to be dissolution, then she will try to take the newly encountered phenomenon to match with this experiment. What can be matched are dissolution, and others are not.

If this conjecture is correct, it will bring considerable enlightenment to the existing science teaching, especially the concept teaching. At present, the primary science education is more advocating the concept construction theory of structuralism, that is, a new learned concept by children should be a complex cognitive structure in their brain. However, the situation of the girl suggests that whether there are many modes of scientific concept learning? And is structuralist model the favorite preference in children’s learning?

The cognitive theory that dominates international primary education has always been structuralist. In 1934, Soviet scholar Lev Vygotsky published the book “Language and Thinking”, in which he proposed the concept learning theory that still has a significant impact on modern science education. He believes that, first of all, children will mix a series of phenomena through observation to form a unique compound: compound thinking. The second step is that children’s brains should process these compounds to form a “chain compound” - the “purest type of compound thinking” (Lev Vigotsky, 2005, P141). Third, children will abstract and summarize these compounds, that is, form a pseudo concept. But this obscure abstraction is extremely unstable and vulnerable to the interference of concrete phenomena. The key to stabilizing it is language. Vygotsky believes that the addition of language to cognitive processes makes pseudo concepts eventually form into scientific concepts. Therefore, Vygotsky’s concept of conceptual learning can be roughly summarized as a process: phenomena → compound thinking → chain compound → pseudo concept → concept (Lev Vygotsky, 2005, P115-181).

In the mid-20th century, with the advent of the Cognitive Revolution, a group of cognitive scientists emerged in the United States that had a profound impact on primary education today. As J. Bruner proposed the theory of “Discovery Learning Theory” (DLT). Compared to traditional learning method as mechanical memory, DLT is achieved through learners’ independent learning and thinking, self-discovering knowledge and mastering principles (Keiichi Takaya, 2015). It can be seen that the theory of DLT has great similarities with Vygotsky’s conceptual construction, therefore ‘as to the cause of development, Bruner is unmistakably Vygotskyan’ (Keiichi Takaya, 2015). If both of the above emphasize that learning is actively constructed by children, then D. Ausubel is clearly more radical than them. He views learning as a child’s active activity and constructs his “Meaningful Learning Theory” (MLT) theory based on learning motivation and past experience. In its view, MLT is a continuous process of interaction between new knowledges and relevant prior knowledges. Therefore, in Ausubel’s view, the most important factor affecting learning is what students previously knew (Glenda Agra et al., 2019).

Unlike the above scholars, B. Bloom shifted his focus from the learning process to the learning objectives, dividing teaching objectives into six levels: Knowledge, Comprehension, Application, Analysis, Synthesis, and Evaluation. In the eyes of scholars after him, Knowledge and Comprehension belong to relatively low order cognitive abilities, while the remaining four are relatively high order abilities (Bloom did not seem to have divided order between low and high order before his death). There are two criteria for dividing these six levels: firstly, to what depth does achieving these criteria mean mastering concepts; the second is the level and difficulty of mobilizing cognitive resources to achieve these levels (Nancy E. Adams, 2015). The influence of Bloom’s theory is extremely profound. If carefully analyzed, it is not difficult to find that although the details are different, the formulation of behavioral goals for standards of science education such as NGSS and large-scale tests such as PISA is based on the perspective of behavioral taxonomy.

The above does not include all influential structuralists (such as J Piaget), we only reviewed the influential people and their views in the international science education community. In the eyes of structuralists, concept learning is a complex and ingenious cognitive architecture that requires systematic processing by the brain. Therefore, teaching from the perspective of this school should involve teachers providing various clues and scaffolding to assist students in fully mobilizing deep cognitive resources through enlightenment, guidance, and other means, thus building a complex cognitive architecture. In other words, learn a new concept by initiative inquiry, and analysis clues in experiments thoroughly is naturally in structuralists’ eye. However, this obviously cannot explain the case at the beginning of this article, as the girl almost without hesitation ascribed to the problem to the prototype she had seen in lesson. This reminds us that there seem to be some inherent preferences and “shortcuts” in children’s concept learning.

Prototype matching is not a new concept in psychology, some previous psychological studies have confirmed that human conceptual learning may follow a variety of ways, and prototype matching is an important one. For example, Rips et al. found that people responded more quickly to the sentence “robin is a bird” than to the sentence “turkey is a bird”. In this regard, K M. Galotti believes that, for most people, robin is a typical bird, but turkey is not (Katheleen M. Galotti, 2017, P113). As Wang Shugen mentioned, “it (prototype matching) believes that the memory stored in the human brain is not a template corresponding to the external model, but a prototype. This model reflects the basic characteristics of a class of objects” (Shugen Wang, 2002). However, it is a pity that at present, the psychological circle does not seem to have much research on the priority of various ways of concept learning, nor does it point out which may be more preferred by children (previous preference research has focused more on the overall learning model, while the organization and preference of concepts and knowledges are relatively rare). Relevant research is more seen in animal experiments. For example, Thomas A. Daniel et al. conducted abstract concept learning training on young pigeons, and found that these pigeons have obvious preference effects (Thomas A. Daniel et al, 2016). However, this paper described pigeons’ preference behavior for matching as “oddity”, indicating that in the eyes of researchers, this concept learning method should not be a representative phenomenon.

In fact, some previous studies have shown that students have shown signs of seeking prototypes in science learning. For example, Baosheng Ye and Xiang Peng found in their research on the pre scientific concept of primary school students that primary school students have the characteristics of “outstanding and obvious characteristics” in concept learning. For example, when they studied the characteristics of children’s textile fabrics, they found that 60% of students believed that the characteristics of textile fabrics were soft. In this regard, the article believes that: “Textile fabrics are things with comprehensive characteristics, but the proportion of students’ understanding of the feature of ‘softness’ is far higher than other features… Pupils only have a clear understanding of their’ softness’, which indicates that they have a distinctive cognitive feature” (Baosheng Ye, Xiang Peng, 2018). Obviously, this “softness” has already been prototype. However, these signs have not attracted the attention of relevant educational scholars, not only have they not been written into the previous curriculum standards, but also can hardly be found in the influential papers on “concept learning”.

## 2. Experimental design

### 2.1 Basic ideas

Because the study is about preference, this experiment adopts distributed fitting as the basic idea. That is, we take “students have no special preference in learning scientific concepts, but only make judgments based on calm analysis” as zero hypothesis H0. And then let the participants (students) complete a series of choice questions (test tasks), all of which have only two options, A and B, one of which contains the prototype they have been exposed to; The other one is not——it requires careful thinking and analysis to determine whether it is correct. According to H0, the test results should be random, so the probability of selecting the option with prototype should be close to 0.5 (when the total number of participants is enough). Finally, we use goodness of fit test to verify whether the actual observed frequency fits 0.5, and then judge whether H0 is acceptable.

### 2.2 Test task design

First of all, because the current knowledge content of biology, universe, earth science in primary school stage almost does not involve experiment and inquiry learning, but use the teaching method based on observation, we can’t screen the students’ preferences through this experiment, so the test task does not include the above content.

Secondly, in order to ensure that the prototype students of the test task have been exposed to it, the prototype options in all topics are based on the experiments (or pictures) in the textbook (Bo Yu, 2020). In order to highlight the experimental effect, we “tampered” some prototypes to make them look like the prototypes they have been exposed to, but after careful analysis, participants can find that they are actually wrong and cannot explain the knowledge points involved in this question.

Third, because there is often many typical experiments or models presented in textbooks, we cannot confirm which of them are the prototype of students, so we designed two sets of questionnaires (test tasks). The knowledge points involved in the two sets of questionnaires are identical, and the non-prototype options are also identical. The only difference is the options include prototypes. See “6. Appendix” for questionnaires. Among them, the red font is the option with prototype, and the question number marked with ^“ *”^ is our option of “tampered”. The red font and ^“ *”^ will not appear in the real questionnaire.

### 2.3 Design of statistical analysis method

We used goodness of fit test as the main statistical method of this study. The reason for choosing this method is that, first, as mentioned above, the research target of this experiment is preference, and second, it can “judge whether there is a significant difference between the expected frequency and the observed frequency” (Baike.baidu, “GOODNESS OF FIT TEST”). Therefore, it can be used to explain whether the deviation between the sample results of this experiment and 0.5 is caused by inevitable random errors.

Previous studies have also shown that goodness of fit test is very suitable for behavioral preference research. For example, Melanie et al. studied the preference of 1282 people in Germany for colorectal cancer screening through this method. Because screening is of great significance for early diagnosis and treatment of colorectal cancer, but it also has certain risks (Baike.baidu, “COLONOSCOPY”), so patients have a game based decision-making problem on whether to choose. Because of that, that paper examined the preference through goodness of fit test (Melanie Brinkmann MSc et al, 2022).

### 2.4 Effect size method design

As mentioned above, this study is based on chi square test (goodness of fit test), so we use Cramer’s V correlation coefficient as the calculation method of the effect size. The calculation formula of this method is:

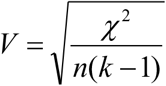

Where, n is the sample size, and k is the smaller of the number of rows and columns (Zheng Haomin, Wen Zhonglin, Wu Yan, 2011).

The size standard of the effect of this method is small effect V∈0.1, medium weak effect V∈(0.1, 0.3], medium effect V∈(0.3, 0.5], large effect V>0.5). (Baike.baidu, EFFECT SIZE)

### 2.5 Sample size design

Since this study is based on goodness of fit test, which objectively requires a high sample size, we should especially avoid conclusion bias caused by too small sample size (the so-called “sparse data”) (Hu Chunyan et al., 2021). For this purpose, we use a large sample size (n>30). At the same time, we require that the number of participants rejected should not exceed 5% (inclusive) of the total number of participants.

## 3. Experiment preparation and implementation

### 3.1 Participants

We selected 160 fifth-grade students from a primary school in Beijing as participants. The reason for choosing the fifth-grade is to ensure that they have learned all the knowledges in the questionnaire and have been exposed to the prototypes. All students have medium academic ability and good IQ. This experiment has passed the ethical review of local institutions, and guaranteed the participant’s right to know to the greatest extent without exposing the purpose of the test.

### 3.2 Instruments

The paper questionnaires were used for data collection in this experiment, SPSS24 software was used for data statistics and analysis, and the analysis instrument was a Redmi Book Pro14 laptop.

### 3.3 Experimental process

This experiment includes two rounds in total. The first round is the test task described above. In the distribution of test papers, the principle of random distribution is adopted, that is, two test papers are randomly mixed together by means of “shuffling” and randomly distributed to the participants by teachers who have not participated in this experiment.

The second round is an additional task, which aims to eliminate a potential competitive explanation: Are the options abandoned by most students those that students have not learned or will not (or even cannot understand)? Although the participants were in the fifth-grade, that is, the knowledge we chose should have been learned by students, because of the epidemic, many students had the experience of studying at home, and objectively it was difficult to guarantee the learning effect, so we added this task. We chose the questions which the participants no selected in the first round of test and redesigned the relatively detailed questions (except for a circuit question, all it was still choice questions) to see whether the students really can do it. In order to ensure that there is no consistency bias in the results (for example, doing the same questions again increases the experience and thinking time of the participants), we chose another class (38 students in total) with the same conditions as the participants. The students in this class have not experienced the first round of test tasks (that is, they haven’t seen these questions), so their answers can reflect the average ability of the participants to deal with the first round of test tasks.

See “6. Appendix” for the second round of test questions (additional tasks).

## 4. Results and Analysis

### 4.1 Data preprocessing

Since most test tasks are options A and B, we must assign values to the answers before statistical analysis. The specific assignment method is as follows: if a participant chooses an option with prototype (no matter whether the option is A or B) when answering a question, we will assign the answer as 1; Otherwise, if the non-prototype option is selected, the value is assigned as 0. The first question in the additional task is to use arrows to indicate the current direction. The correct answer should be 6 arrows. We calculate the number of arrows drawn by the participants and divide it by 6 to get the assignment of the question (if a participant draws 4 arrows correctly, we will assign 4 ÷ 6 ≈ 0.67). Since the positive and negative poles of the battery cannot be seen clearly, the current direction can be ignored, but all arrows drawn should follow the same direction. This assignment ensures that the data points to the point of “whether the probability of selecting the prototype is random”, which is the focus of this study.

At the same time, we eliminated the answers of 5 participants, mainly because of missed questions or multiple choices. The number of rejected products accounted for 3.13% of the total, which was in line with the sample size design (see 2.5 in this article for details).

The data after preprocessing are as follows:

**Table 1:**
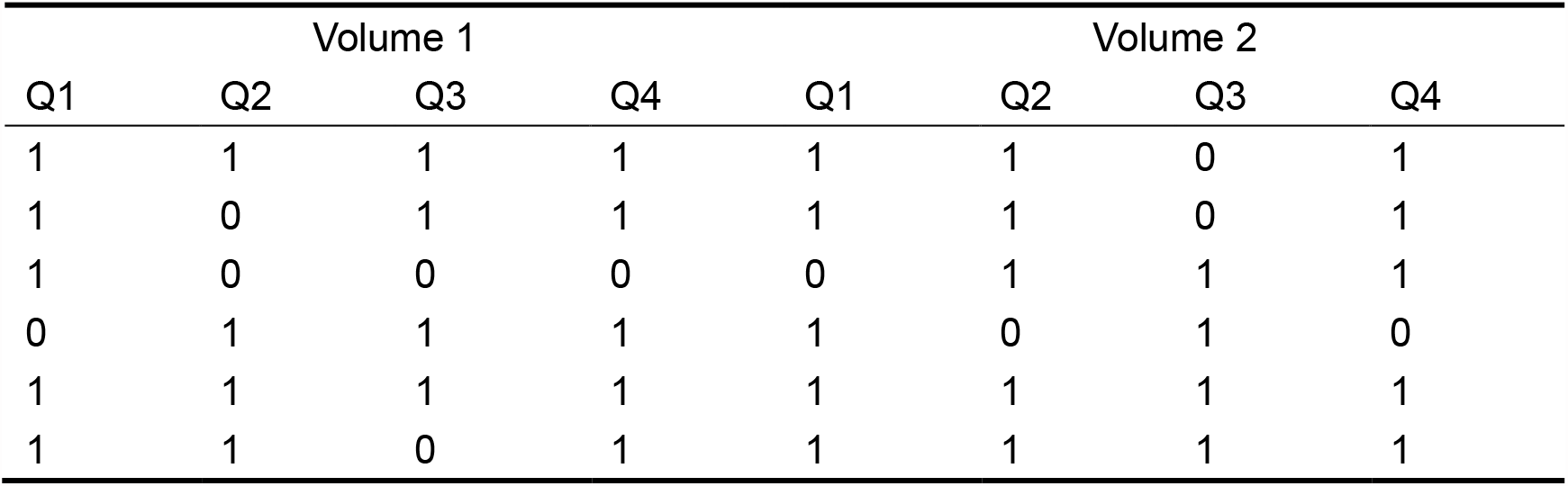
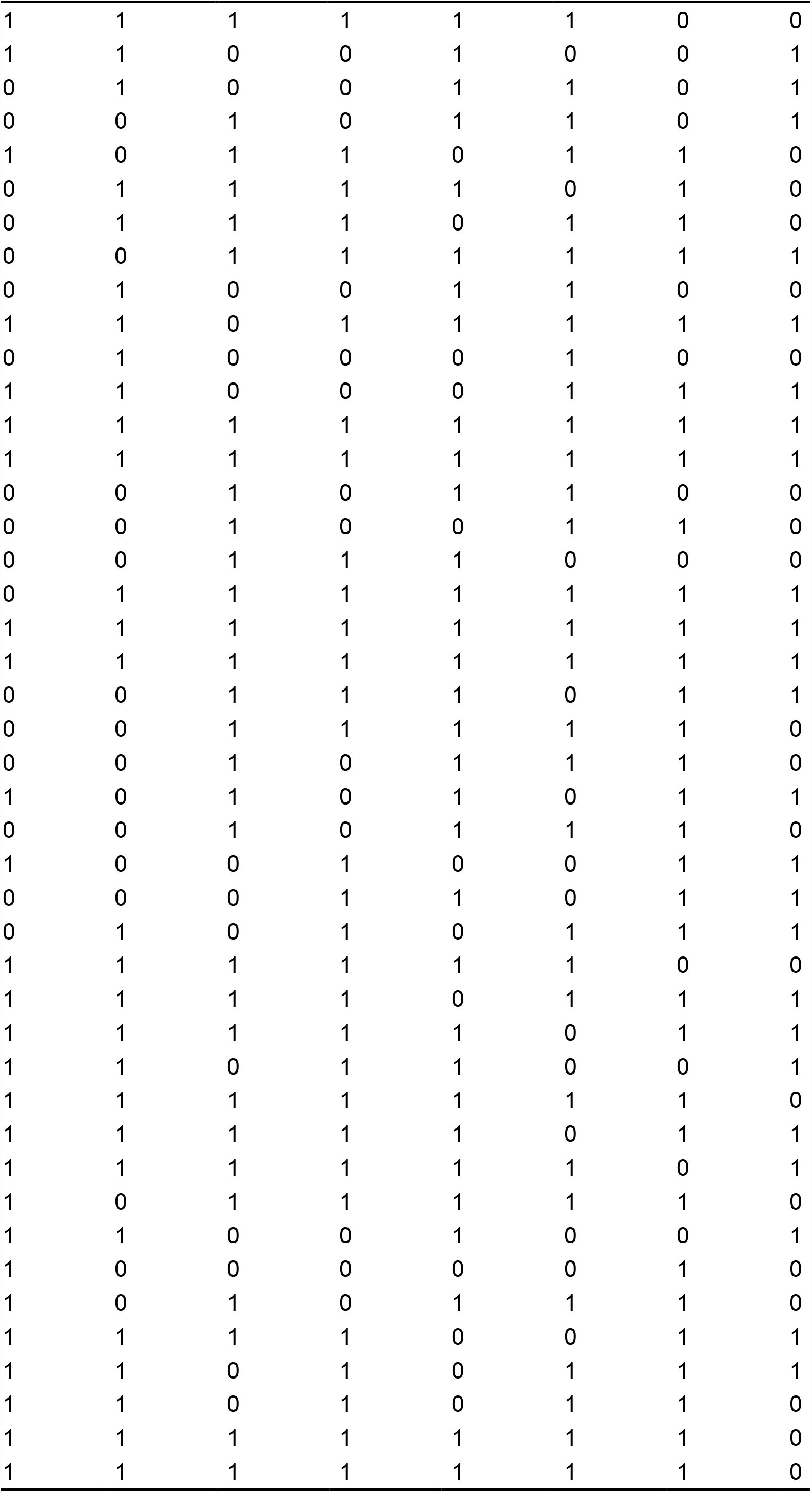
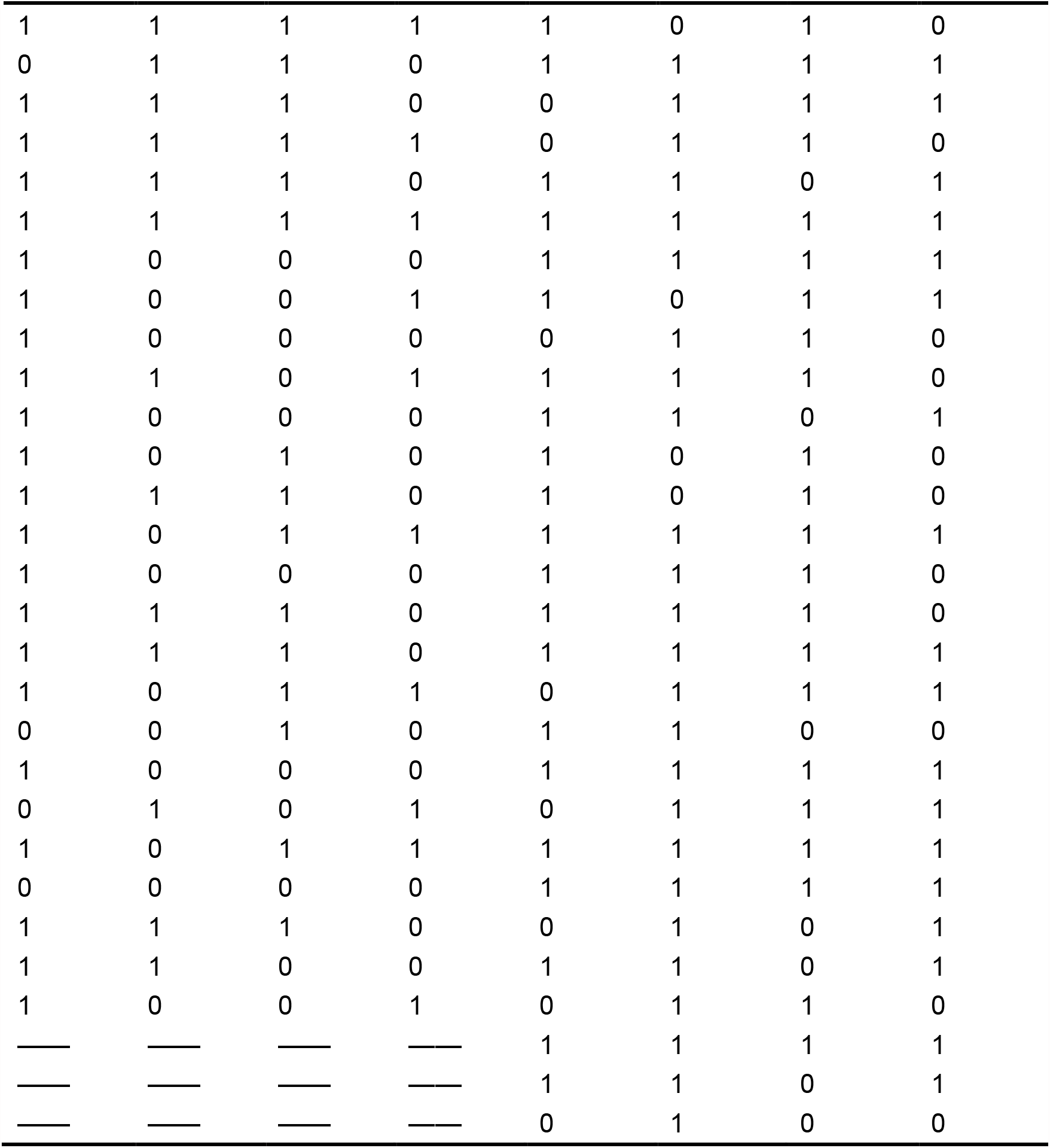
Test Task Results.

| Volume 1 |  |  |  | Volume 2 |  |  |  |
| --- | --- | --- | --- | --- | --- | --- | --- |
| Q1 | Q2 | Q3 | Q4 | Q1 | Q2 | Q3 | Q4 |
| 1 | 1 | 1 | 1 | 1 | 1 | 0 | 1 |
| 1 | 0 | 1 | 1 | 1 | 1 | 0 | 1 |
| 1 | 0 | 0 | 0 | 0 | 1 | 1 | 1 |
| 0 | 1 | 1 | 1 | 1 | 0 | 1 | 0 |
| 1 | 1 | 1 | 1 | 1 | 1 | 1 | 1 |
| 1 | 1 | 0 | 1 | 1 | 1 | 1 | 1 |
| 1 | 1 | 1 | 1 | 1 | 1 | 0 | 0 |
| 1 | 1 | 0 | 0 | 1 | 0 | 0 | 1 |
| 0 | 1 | 0 | 0 | 1 | 1 | 0 | 1 |
| 0 | 0 | 1 | 0 | 1 | 1 | 0 | 1 |
| 1 | 0 | 1 | 1 | 0 | 1 | 1 | 0 |
| 0 | 1 | 1 | 1 | 1 | 0 | 1 | 0 |
| 0 | 1 | 1 | 1 | 0 | 1 | 1 | 0 |
| 0 | 0 | 1 | 1 | 1 | 1 | 1 | 1 |
| 0 | 1 | 0 | 0 | 1 | 1 | 0 | 0 |
| 1 | 1 | 0 | 1 | 1 | 1 | 1 | 1 |
| 0 | 1 | 0 | 0 | 0 | 1 | 0 | 0 |
| 1 | 1 | 0 | 0 | 0 | 1 | 1 | 1 |
| 1 | 1 | 1 | 1 | 1 | 1 | 1 | 1 |
| 1 | 1 | 1 | 1 | 1 | 1 | 1 | 1 |
| 0 | 0 | 1 | 0 | 1 | 1 | 0 | 0 |
| 0 | 0 | 1 | 0 | 0 | 1 | 1 | 0 |
| 0 | 0 | 1 | 1 | 1 | 0 | 0 | 0 |
| 0 | 1 | 1 | 1 | 1 | 1 | 1 | 1 |
| 1 | 1 | 1 | 1 | 1 | 1 | 1 | 1 |
| 1 | 1 | 1 | 1 | 1 | 1 | 1 | 1 |
| 0 | 0 | 1 | 1 | 1 | 0 | 1 | 1 |
| 0 | 0 | 1 | 1 | 1 | 1 | 1 | 0 |
| 0 | 0 | 1 | 0 | 1 | 1 | 1 | 0 |
| 1 | 0 | 1 | 0 | 1 | 0 | 1 | 1 |
| 0 | 0 | 1 | 0 | 1 | 1 | 1 | 0 |
| 1 | 0 | 0 | 1 | 0 | 0 | 1 | 1 |
| 0 | 0 | 0 | 1 | 1 | 0 | 1 | 1 |
| 0 | 1 | 0 | 1 | 0 | 1 | 1 | 1 |
| 1 | 1 | 1 | 1 | 1 | 1 | 0 | 0 |
| 1 | 1 | 1 | 1 | 0 | 1 | 1 | 1 |
| 1 | 1 | 1 | 1 | 1 | 0 | 1 | 1 |
| 1 | 1 | 0 | 1 | 1 | 0 | 0 | 1 |
| 1 | 1 | 1 | 1 | 1 | 1 | 1 | 0 |
| 1 | 1 | 1 | 1 | 1 | 0 | 1 | 1 |
| 1 | 1 | 1 | 1 | 1 | 1 | 0 | 1 |
| 1 | 0 | 1 | 1 | 1 | 1 | 1 | 0 |
| 1 | 1 | 0 | 0 | 1 | 0 | 0 | 1 |
| 1 | 0 | 0 | 0 | 0 | 0 | 1 | 0 |
| 1 | 0 | 1 | 0 | 1 | 1 | 1 | 0 |
| 1 | 1 | 1 | 1 | 0 | 0 | 1 | 1 |
| 1 | 1 | 0 | 1 | 0 | 1 | 1 | 1 |
| 1 | 1 | 0 | 1 | 0 | 1 | 1 | 0 |
| 1 | 1 | 1 | 1 | 1 | 1 | 1 | 0 |
| 1 | 1 | 1 | 1 | 1 | 1 | 1 | 0 |

*Table 1: Test Task Results*
|  |  |  |  |  |  |  |  |
|---|---|---|---|---|---|---|---|
| 1 | 1 | 1 | 1 | 1 | 0 | 1 | 0 |
| 0 | 1 | 1 | 0 | 1 | 1 | 1 | 1 |
| 1 | 1 | 1 | 0 | 0 | 1 | 1 | 1 |
| 1 | 1 | 1 | 1 | 0 | 1 | 1 | 0 |
| 1 | 1 | 1 | 0 | 1 | 1 | 0 | 1 |
| 1 | 1 | 1 | 1 | 1 | 1 | 1 | 1 |
| 1 | 0 | 0 | 0 | 1 | 1 | 1 | 1 |
| 1 | 0 | 0 | 1 | 1 | 0 | 1 | 1 |
| 1 | 0 | 0 | 0 | 0 | 1 | 1 | 0 |
| 1 | 1 | 0 | 1 | 1 | 1 | 1 | 0 |
| 1 | 0 | 0 | 0 | 1 | 1 | 0 | 1 |
| 1 | 0 | 1 | 0 | 1 | 0 | 1 | 0 |
| 1 | 1 | 1 | 0 | 1 | 0 | 1 | 0 |
| 1 | 0 | 1 | 1 | 1 | 1 | 1 | 1 |
| 1 | 0 | 0 | 0 | 1 | 1 | 1 | 0 |
| 1 | 1 | 1 | 0 | 1 | 1 | 1 | 0 |
| 1 | 1 | 1 | 0 | 1 | 1 | 1 | 1 |
| 1 | 0 | 1 | 1 | 0 | 1 | 1 | 1 |
| 0 | 0 | 1 | 0 | 1 | 1 | 0 | 0 |
| 1 | 0 | 0 | 0 | 1 | 1 | 1 | 1 |
| 0 | 1 | 0 | 1 | 0 | 1 | 1 | 1 |
| 1 | 0 | 1 | 1 | 1 | 1 | 1 | 1 |
| 0 | 0 | 0 | 0 | 1 | 1 | 1 | 1 |
| 1 | 1 | 1 | 0 | 0 | 1 | 0 | 1 |
| 1 | 1 | 0 | 0 | 1 | 1 | 0 | 1 |
| 1 | 0 | 0 | 1 | 0 | 1 | 1 | 0 |
| — | — | — | — | 1 | 1 | 1 | 1 |
| — | — | — | — | 1 | 1 | 0 | 1 |
| — | — | — | — | 0 | 1 | 0 | 0 |

**Table 2:**
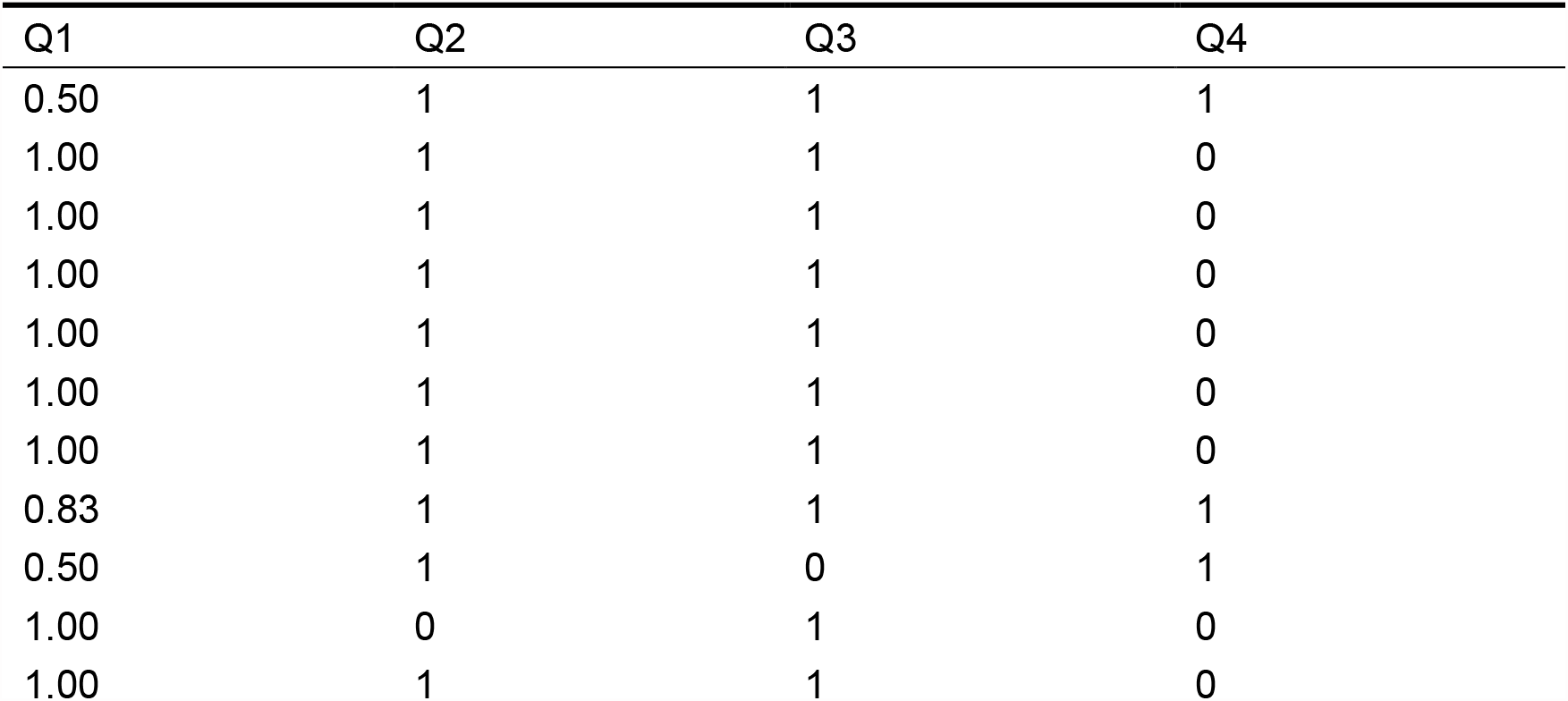
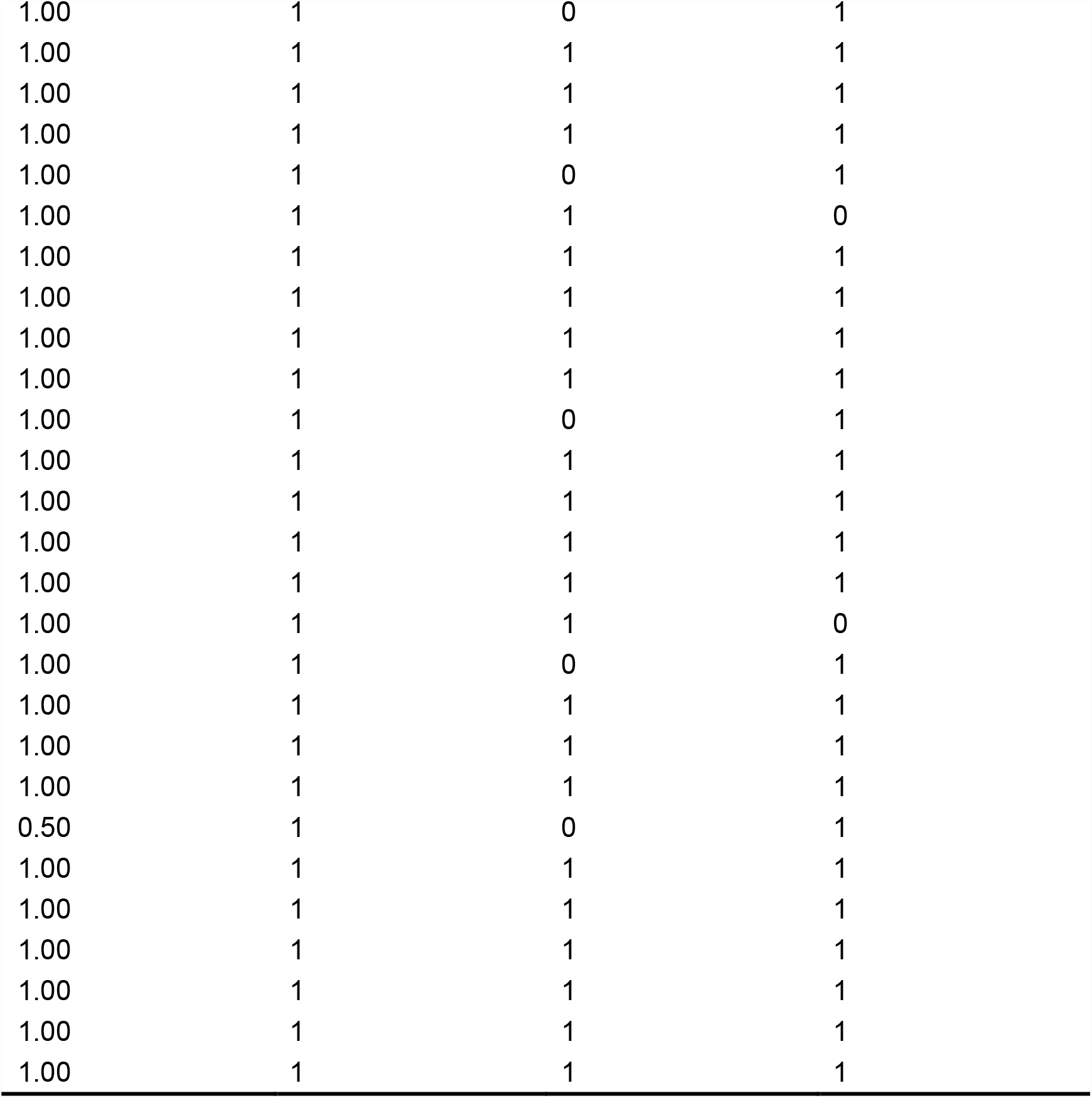
Results of additional tasks.

### 4.2 Excluding Competitive Interpretation

As described in 3.3 of this article, in order to prevent competitive interpretation - that is, the reason why these options are not selected is likely to be because we cannot do/understand the topic - we added a round of tests. The main content of this round of test is to select those options that are less selected by the participants in the test task separately, and make a separate proposition to see whether the students with average learning ability of the participants can answer these questions.

Table 3 shows the accuracy of 38 students’ additional tasks (rounded to two decimal places). It can be seen that the average correct rate of students with the same academic level is 0.88, that is, as long as they think carefully, they will completely make these choices. Therefore, the competitive interpretation is basically excluded.

**Table 3:**
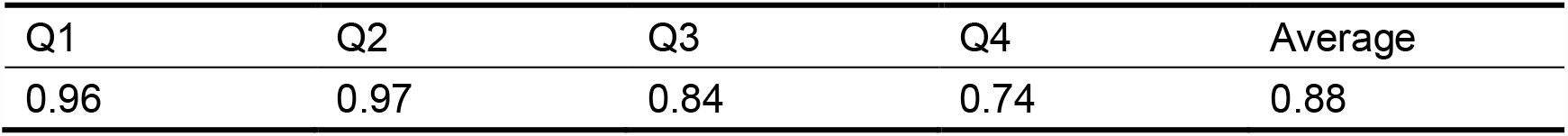

### 4.3 Inspection results and analysis

#### 4.3.1 Overall statistical results

Table 4-6 shows the results after goodness of fit test, of which Table 4 is the descriptive statistics. It can be seen that the chi square value of the overall test task is 73.865, with a significance of P<0.001, and there is no cell with the expected frequency lower than 5. The correlation coefficient of Cramer’s V, V ≈ 0.35, has a medium effect. It is easy to see that the participants have obvious prototype preference in their choice, and this preference has very obvious statistical significance.

**Table 4:**

| Cases | Average | SD | Min | Max |
| --- | --- | --- | --- | --- |
| 620 | 0.6726 | 0.46965 | 0.00 | 1.00 |

**Table 5:**
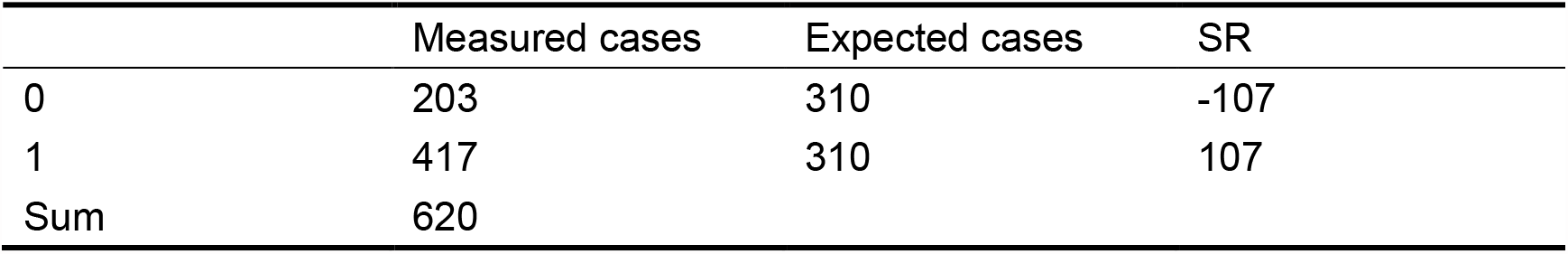

**Table 6:**
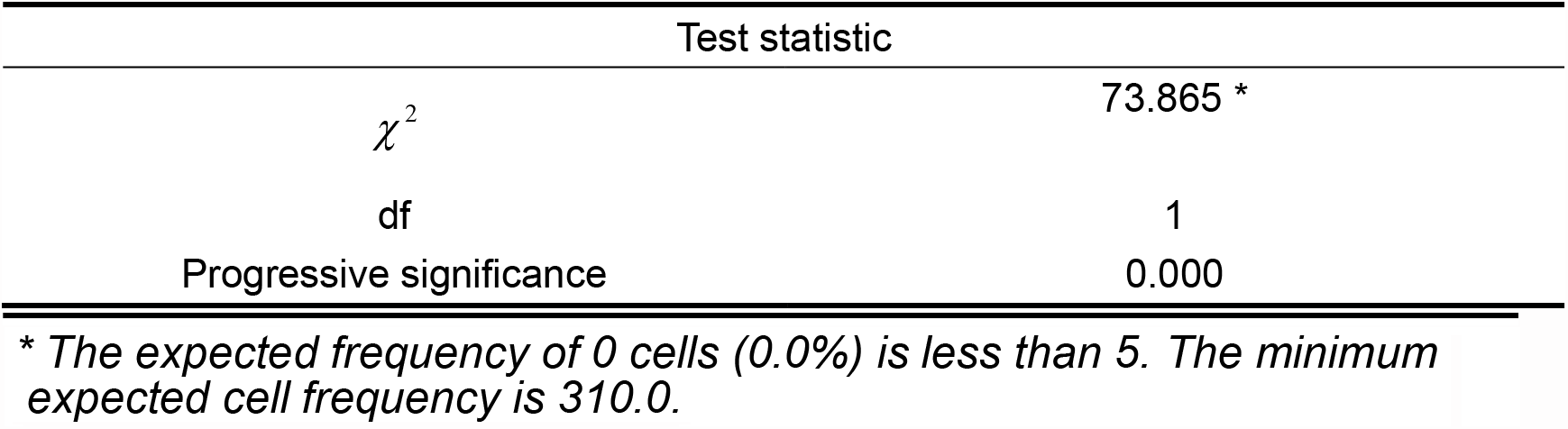

| Test statistic |  |
| --- | --- |
| $\chi^2$ | 73.865 * |
| df | 1 |
| Progressive significance | 0.000 |

\* The expected frequency of 0 cells (0.0%) is less than 5. The minimum expected cell frequency is 310.0.

#### 4.3.2 The questions which “be tampered”

Although it can be seen from the data that the participants chose many prototype options, is it because those options are also correct? For this reason, we specially select the statistical results of those “tampered” topics with the “prototype” option (that is, the so-called “prototype” is actually wrong and cannot explain the scientific phenomenon referred to in the topic). Table 7-9 shows the statistical results of those topics. The effect size of Cramer’s V≈0.31, has a medium effect. Similar to the approximate results shown in Table 4-6, the participants still had obvious prototype preference. More importantly, although there are pitfalls in these questions, it seems that the participants did not pay attention to this.

**Table 7:**
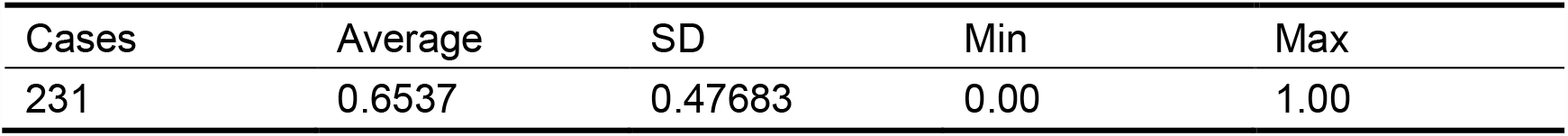

**Table 8:**

|  | Measured cases | Expected cases | SR |
| --- | --- | --- | --- |
| 0 | 80 | 115.5 | -35 |
| 1 | 151 | 115.5 | 35 |
| Sum | 231 |  |  |

**Table 9:**

| Test statistic |  |
| --- | --- |
| $\chi^2$ | 21.823 * |
| df | 1 |
| Progressive significance | 0.000 |

\* The expected frequency of 0 cells (0.0%) is less than 5. The minimum expected cell frequency is 115.5.

#### 4.3.3 Behavioral difference of whether “be tampered”

What’s more, do the participants have obvious behavioral differences in answering the two types of questions, “Have you ever done anything?”? If they exist, it is likely that these problems affect the students (of course, it also means that the students are aware of these tricks), otherwise, they do not. In this regard, we used independent sample t-test to examine the differences of the participants in answering the two questions.

**Table 10:**

| | Cases | Average | SD | $\sigma$ |
| --- | --- | --- | --- | --- |
| "Be tampered" | 389 | 0.6838 | 0.46559 | 0.02361 |
| Never "be tampered" | 231 | 0.6537 | 0.47683 | 0.03137 |

**Table 11:**
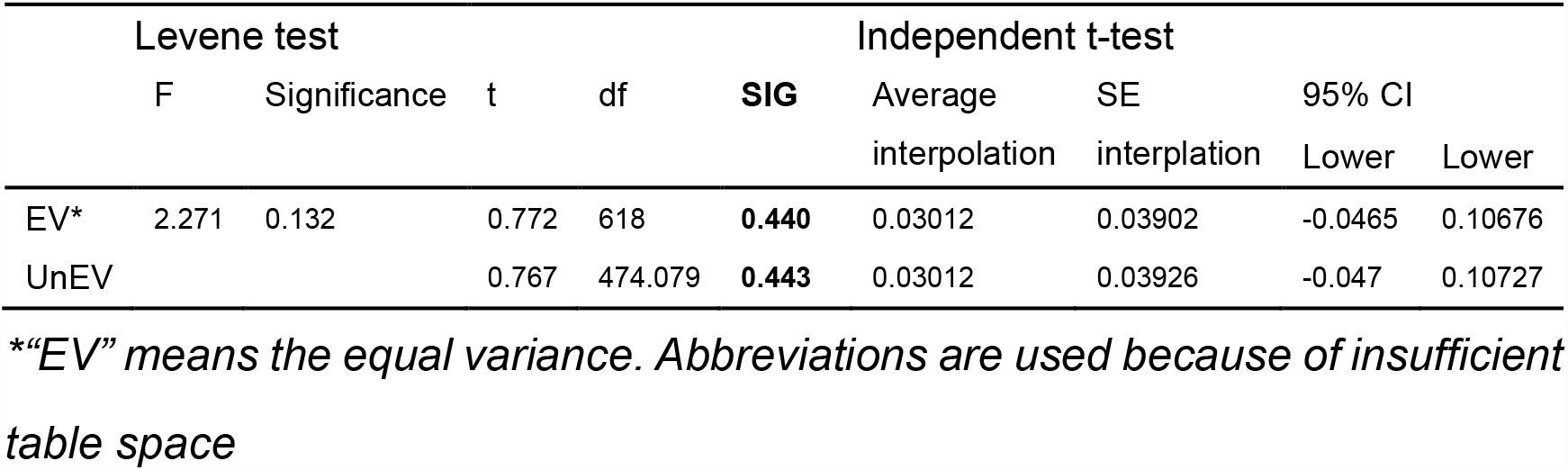

It can be seen from the above table that there is no significant difference in the data, which indicates that the participants have no evidence to prove that they are aware of these tricks when answering the question of whether they have “be tampered”. From the data, it seems that the participants are likely to have similar behaviors on these two types of questions.

To sum up the data analysis, we believe that the zero hypothesis H0 can basically be denied, that is, from the data point of view, students have an obvious preference for prototype matching when identifying and judging concepts. More persuasively, even though the so-called “archetypes” were wrong, the participants still stubbornly chose them. This seems to reflect that the participants’ choice of prototype is instantaneous and intuitive, because it is not difficult to find these traps as long as they think a little.

## 5. Discussion

This article tested the preference of 160 pupils for learning scientific concepts. The results show that, unlike mainstream structuralists who believe that the process of concept learning should be actively constructing a complex cognitive structure, children prefer to learn new concepts through a shortcut as prototype matching. And results shows that this preference seems to be automatic, that is, there is no evidence of students making judgments by comparing with the definition.

Concept learning through prototypes is a well-established method of concept learning (Dagmar Zeithamova et al, 2019), and relevant evidence shows that humans prefer this learning method when learning concrete concepts (Gabriel Recchia et al, 2012). Specifically, the working mechanism of this learning approach seems to rely on shared features between concepts (old knowledges learned and the new concept that need to be learned) (Gabriel Recchia et al, 2012). For example, an apple is a typical fruit that has many common characteristics (seeds, skin, sweetness, and juiciness), while an avocado have more characteristics that fruits do not have (saltiness, high fat content), making it difficult to associate avocados with fruits (Anna M. Woollams, 2012). Therefore, concepts with more obvious shared features are more likely to be identified as the same class and activated together. The reason why the phrase ‘more obvious’ is used here is because there is still controversy about what characteristics of shared features cause activation, with some evidence leaning towards quantity (Gabriel Recchia et al, 2012) and others towards feature correlation (Anna M. Woollams, 2012).

This article demonstrated from the perspective of science education that the learning of primary school scientific concepts is more easily accomplished through concept sharing features (especially prototype matching). To our knowledge, although the “prototype” is not a new concept in cognition and neuroscience, few people have conducted such empirical experiments in current primary education, to prove the role of prototype in concept learning. Considering that most concepts in primary education (especially science education) are concrete, and there are not many abstract concepts such as “energy” and “ecological environment”, it is reasonable to believe that prototype matching is one of the most important ways for children to learn scientific concepts.

More interestingly, this article seems to provide side evidence for a long-standing issue: competition and attention allocation in semantic processing. Competition in semantic processing is a common phenomenon, which not only includes the interpretation of competitive interpretation of ambiguities parts in the same sentence (Jennifer Rodd et al, 2002), but also includes competition between visual attention. A more convincing explanation for this issue is that the amount (and difficulty) of brain resources mobilized by both competing parties determines the outcome of the competition, such as Kleinman D believes that in image vocabulary interference tasks, the cognitive resources mobilized by images and vocabulary greatly affect the results of such stimuli (Kleinman D, 2013). For this study, why do students choose that “tampered” prototype options without careful consideration? Using the above explanation makes it easier to understand students’ behavior: activating a prototype requires less cognitive resources than proving it, so activating a prototype is clearly more competitive than understanding and thinking word for word. The learning and activation of prototypes is considered one of the most convenient and efficient classification methods, and has even been proven in many animals (such as pigeons, jackdaws, and primates) (Apostel, A. et al, 2023). In other words, students’ unconscious choice of that “tampered” prototype options may indirectly prove that the difficulty of cognitive processing is one of the determining factors in attention competition.

If the above inference is credible, it not only indicates that prototype matching is the main way for children to learn concrete concepts, but also means that it occupies a priority position among many concept processing methods. The significance of this for current primary education is self-evident: the development of scientific education standards and large-scale testing goals should not only consider the requirements of discipline teaching, but also pay more attention to children’s cognitive preferences and characteristics.

## 8. Full text conclusion

First, children have a clear preference for “prototype matching” in scientific concept learning, which is not only a clear deviation from the current general understanding of science education, which emphasizes discovery/inquiry construction, but also points out that various ways of concept organization (such as definition theory, prototype theory, schema theory, etc.) may have a priority relationship. Second, children’s preference for archetypes seems to be unthinking, and they will not identify the authenticity of archetypes, which is particularly noteworthy in first-line teaching.

## 6. Appendix

All experimental data, test questions (including additional tasks) have been uploaded to the Scientific Data Bank (ScienceDB). Please click the link to download or view them: https://www.scidb.cn/en/detail?dataSetId=e3bddcd1754b422ba5870fef94346c13

Please click the link to view the interview video of the girl about the case at beginning of this article: http://www.bdice.ac.cn/specimens.html?newsid=291548&_t=1666760039

## Notes

https://www.scidb.cn/en/detail?dataSetId=e3bddcd1754b422ba5870fef94346c13

http://www.bdice.ac.cn/specimens.html?newsid=291548&_t=1666760039

### Competing Interest Statement

The authors have declared no competing interest.

### Summary of Updates

We have revised the introduction and discussion sections to enrich some of the literature.

